# Bioluminescent Multi-Characteristic Opsin for monitoring visual cortical activity upon optical stimulation

**DOI:** 10.1101/509323

**Authors:** Subrata Batabyal, Sivakumar Gajjeraman, Takeharu Nagai, Weldon Wright, Samarendra Mohanty

## Abstract

Non-invasive detection of neural activity is important for the diagnosis of neurological diseases, evaluation of therapeutic outcomes, and collection of real-time feedback for stimulation based therapeutic approaches. In the case of vision loss, due to retinal degeneration, optic neuropathy, or enucleation of the eye, there is a need to map changes in visual cortical activity during disease progression and subsequent vision restoration by retinal, optic nerve or cortical stimulation. Existing technologies allow interrogation of neuronal circuits by both read and write activity, albeit with inherent limitations. Here, we present the development of bioluminescent multi-characteristic opsin (bMCO-11), which is comprised of a highly photosensitive ambient light-activatable domain and a Ca^2+^-sensitive bioluminescence reporter. The high quantum efficiency of bMCO-11 enables light activation and recording of cellular activity upon local as well as wide area optical stimulation. Furthermore, persistent Ca^2+^ influx was achieved by bioluminescence based cyclic activation of the opsin-domain of bMCO-11 by transporting the ions only into bMCO expressing cells. This allowed us to continuously monitor visual cortical activity in wild type and retinal degenerated mice, without requiring any additional external excitation source.

## INTRODUCTION

Dysfunction of the retina, due to photoreceptor degeneration (as in Age-related macular degeneration[1-3] or Retinitis Pigmentosa[4,5]), optic nerve damage (due to Glaucoma or diabetic retinopathy), or enucleation of the eye (by combat ocular trauma[6] or traumatic optic neuropathy[7,8]), leads to a loss of signal transduction and/or transmission to the visual cortex. The electrical and optical stimulation methods employed in clinical trials of vision augmentation in blind patients with degenerated retinas [with intact optic nerves), involve retinal implants or external optical stimulation[9]. Recent optogenetic studies in retinal degenerated mouse models suggest that the photoreceptor-degenerated retina can be re-photosensitized, resulting in vision enhancement at ambient light levels without requiring an external stimulation source[9]. Further, in the case of severely damaged eyes or injured optic nerves, significant attempts have been made to electricalyl stimulate the primary visual cortex[10,11]. To monitor disease progression, and subsequent vision restoration by retinal, optic nerve, or cortical stimulation[12], there is a need to develop safe minimally invasive tools to modulate and map neural activity in the retina and visual cortex.

Current methods for non-invasive detection of visual system activity include electrical and optical functional imaging. While conventional electroencephalography (EEG) and electroretinogram [ERG] [13,14] electrodes monitor visual cortex/retinal activity from outside the skull or eye, electrocorticography [ECoG] [15,16] monitors visual cortex electrophysiological activity by directly placing electrodes on the exposed surface of the brain. These electrode-based detection methods have limited spatial resolution and are prone to cross-talk between their electrical stimulation channels. The use of calcium dyes or genetically encoded voltage/calcium indicators[17-20] have drawbacks, such as cytotoxicity and photobleaching, since observation requires continuous excitation illumination[21]. Although fluorescence imaging is widely used for neural activity monitoring, its dependence on external excitation light induces phototoxicity[21-25] and can hinder longer time-frame observations of neural activity in living organisms. Additionally, since photoreceptors in the retina are also excited by fluorescence excitation light, it is significantly difficult to decouple the signal contribution of visual excitation lignt induced neural activity from unwanted fluorescence excitation light induced neural activity. Moreover, retinal and cortical tissues are auto-fluorescent, which creates a high background signal when collecting fluorescence-based measurements. When using simultaneous optical/optogenetic stimulation, further difficulties arise during fluorescence data analysis when decoupling activation light from external excitation light.

We addressed these difficulties by fusing an ambient light sensitive opsin [MCO-11] with a bright Ca^2+^-sensing bioluminescent protein: *Nano-Lantern-*GenL. Here, we report the development of bMCO-11, a chimeric bioluminescent opsin, which has multiple characteristics of optical detection [read] and activation [write] of neural activity. Using bandwidth engineering, we have synthesized bioluminescent MCO [bMCO-11] to have a broad activation spectrum. Sensitization of the visual cortex with the chimeric bMCO-11 enabled the comprehensive interrogation of visual cortical activity in wild type and retinal degenerated mice, which opens up the capability to concurrently read and write visual system activity. Inclusion of the calcium/voltage sensitive bioluminescent probe (GeNL-Ca^2+^) in this chimeric construct potentially allows for the simultaneous activation and recording of neural activity upon illumination by spectrally separated light corresponding to the opsin activation spectrum.

## RESULTS

### BMCO-11 is stably expressed in cells

In Fig. 1A, we show a typical circular plasmid map showing the insertion of the bMCO-11 gene cloned at the restriction sites (XhoI and BamHI). The putative three-dimensional arrangement of the 1060 amino acid residue protein chain of bMCO-11 (Fig. 1A) was obtained using theoretical modeling by *RaptorX*. Furthermore, the secondary structure prediction [Suppl. Fig. 1] confirms that the bMCO-11 protein consists of 39% α-helix, 22% β-sheet and 37% coiled structure. The DNA gel electrophoresis of bMCO-11 plasmid after BamH/SalI restriction digest is shown in Fig. 1B. HEK293 cells were transfected with bMCO-11 by lipofection and imaged via confocal fluorescence microscopy. In Fig. 1C, the fluorescence image of bMCO-11 transfected HEK293 cells is shown.

**Fig. 1.**
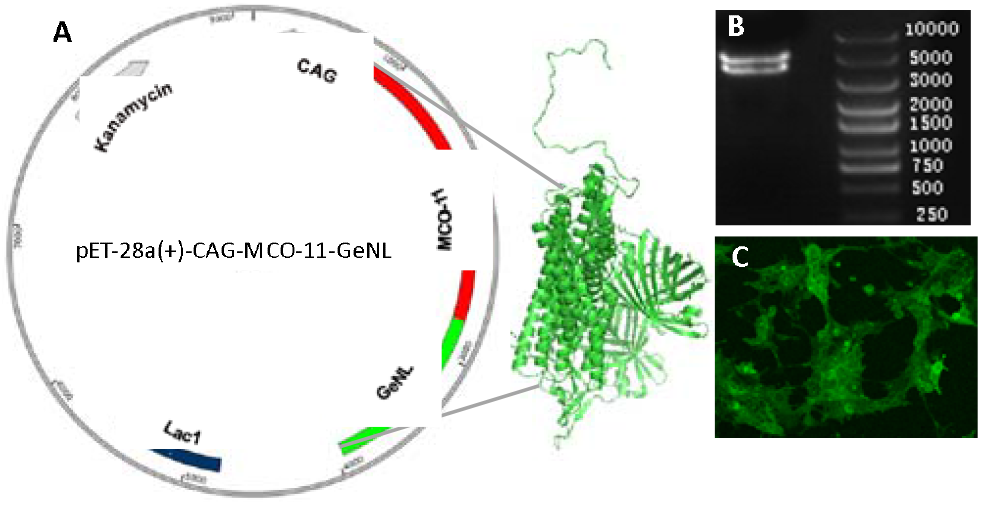
Bioluminescent Multi-Characteristic Opsin (bMCO-11). (A) Vector map of bMCO-11. (B) DNA gel electrophoresis of bMCO-11 plasmid after BamH/SalI restriction digest (Top band: Vector, Bottom band: Gene). (C) Fluorescence image of the bMCO-11 transfected HEK293 cells.

### Bioluminescent Multi-Characteristic Opsin enabled localized optical stimulation and activity detection at single cell level

Expression of chimeric bioluminescent Multi-characteristic opsin [bMCO-11] in HEK293 cells is shown in the confocal fluorescence image in Fig. 2A. A zoomed fluorescence image of cells expressing bMCO-11 is shown in Fig. 2B. Upon localized light stimulation of a targeted bMCO-11 expressing cell, bioluminescence signal increased significantly compared to that of the non-stimulated cells. Suppl. Fig. 2 reveals the kinetics of the changes in the bioluminescence signal [ΔI/I_0_] of the optically stimulated cell in contrast to the control. Though the kinetics of the Ca^2+^ sensing domain [GeNL] of bMCO-11 is known to have sub-sec kinetics, the long-lasting bioluminescence observed upon stimulation of bMCO-11 expressing cells suggests that the bioluminescence light [generated due to short-term activation of the light-activatable domain of bMCO-11] up-regulates the transport of ions [including Ca^2+^], which in turn activates the targeted cell in a cyclic manner.

**Fig. 2.**
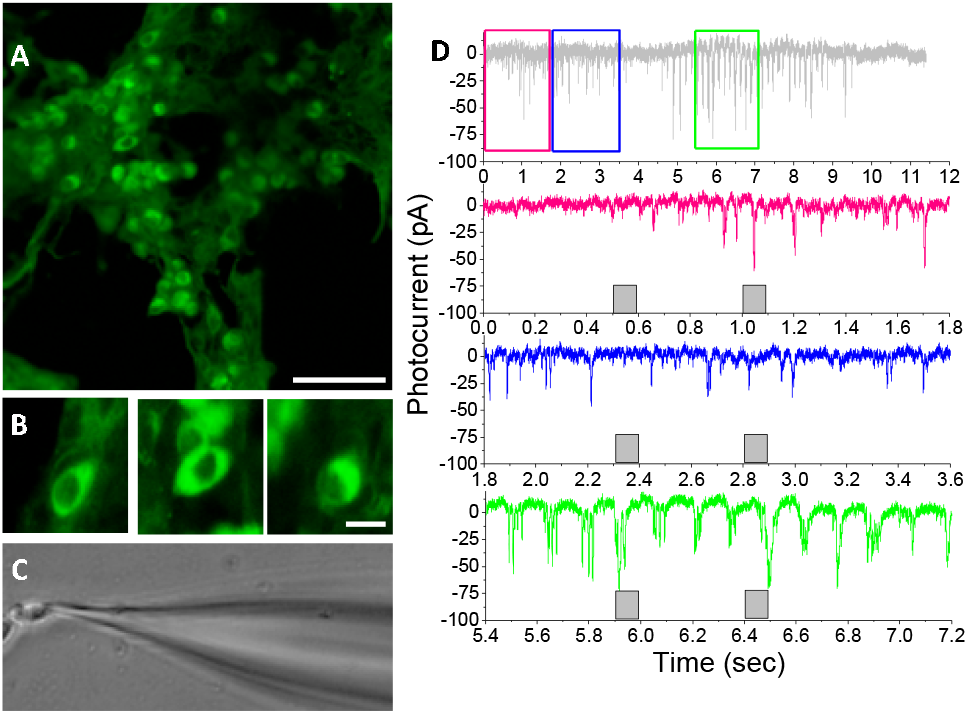
Localized optical stimulation and activity detection at single cell level using Bioluminescent Multi-Characteristic Opsin (bMCO-11). (A) Fluorescence image of HEK 293 cells showing expression of bMCO-11, Scale bar: 30 μm. (B) Zoomed fluorescence images of cells showing membrane expression of bMCO-11. (C) Bright field image of a transfected HEK cell being patched by a glass electrode. (D) Electrophysiology of bMCO-11 sensitized HEK cell showing cyclic activation of opsin-domain by bioluminescence emitted after initial photo-induced current, leading to inward current flows lasting >10 sec, measured by patch-clamp electrophysiology. Optical stimulation light intensity: 37 μW/mm¿.

### Long-lasting activity of bMCO-11 transfected cells observed in electrophysiological measurements

Cyclic activation of the opsin-domain by bioluminescence emitted after the initial light stimulation, was observed to generate inward current flows lasting >10 sec, measured by voltage-clamp electrophysiology. Fig. 2D shows sequential profiles of inward currents in bMCO-11 expressing cells (in presence of ATR and coelenterazine-h), which lasted even after switching off the optical stimulation pulses (100 ms). The minimum light intensity (i.e. threshold) necessary to activate the bMCO-11 expressing cells was found to be 37 μW/mm^2^. The bioluminescence-evoked depolarization generated by bMCO-11 could modulate the intrinsic excitability of the transfected HEK cells in the presence of Coelenterazine giving rise to short transient inwards currents. The kinetics of the short-transient nature of the individual inward current suggests spontaneous fluctuations of membrane current and Ca^2+^ influx by bioluminescence emitted from the GeNL domain.

### Synchronized activation of bMCO-11 expressing cells monitored by bioluminescence upon wide-area repeated light stimulation

Wide-area (1 mm × 1mm) optical stimulation of HEK cells expressing bMCO-11 allowed us to screen the activity of the cells in a high throughput manner (Fig. 3) by simultaneously measuring the bioluminescence activity of a large number of cells. Figs. 3a to c show the time-lapse bioluminescence mapping of cellular activity after light stimulation in the presence of coelenterazine-h[26]. Over 90% of bMCO-11 expressing cells were found to be activated (indicated by a rise in bioluminescence Ca^2+^ signal) in a synchronized manner upon light stimulation. The cells could be repeatedly stimulated and monitored using this approach (Figs. 3d to i). Fig. 3j shows zoomed Ca^2+^ bioluminescence images of the HEK cells. The kinetics of bioluminescence in different cells in response to repeated light stimulation pulses (intensity: 22 μW/mm^2^) is shown in Fig. 3k. The decay in the maximal signal generated during repeated stimulation is attributed to the known instability of the substrate (coelenterazine-h).

**Fig. 3.**
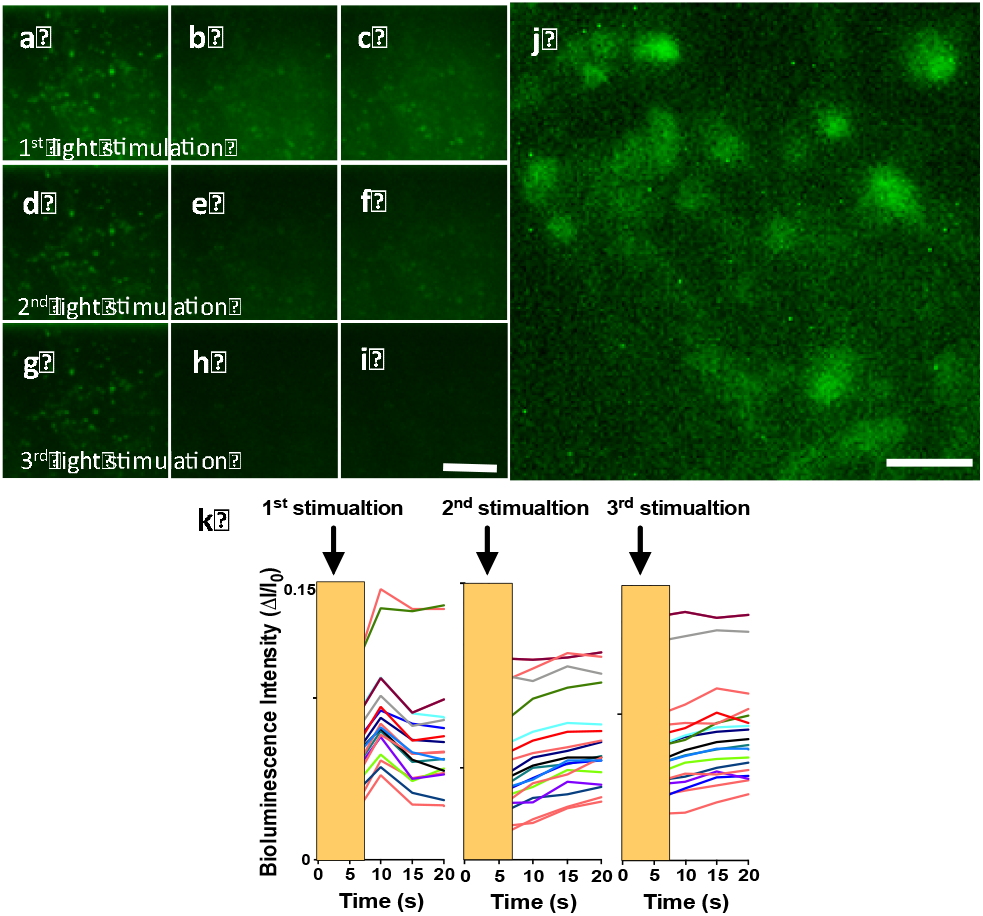
Wide-area light stimulated activity in HEK293 cells expressing bMCO-11 monitored by Ca^2+^-dependent bioluminescence. Bioluminescence mapping at 1,10 and IS seconds after 1^st^ (a-c), 2^nd^ (d-β and 3^rd^ (g-i) light stimulation (marked by arrow in j). Scale bar: 100 μm. (¡) Zoomed image of HEK bioluminescence after stimulation. Scale bar: 20 μm. (k) Kinetics of bioluminescence in different cells in response to light stimulation (intensity: 22 μW/mm^2^).

### Sustained visually evoked bioluminescence activity observed in visual cortices of wild type mice

bMCO-11 expression in the visual cortex was confirmed by epifluorescence imaging [Fig. 4C] via an optical window [Fig. 4B]. Further, serial-sectioned confocal fluorescence microscopic images [Suppl. Fig. 3] showed widespread cellular expression in the primary visual cortex. Fig. 4A shows the schematic of the setup for visual stimulation and bioluminescence imaging in the visual cortex. Time-lapse imaging and kinetics (Fig. 4D) of the bioluminescence signal in dark-adapted (and pupil dilated] wild type mice V1 (with no bMCO-11 transfection and no visual stimulation light) shows flat baseline activity. No change in V1-bioluminescence in the negative control (no bMCO-11) mice was observed even with visual stimulation at an intensity of 20 μW/mm^2^ near eye [Fig. 4D]. In contrast, the mice with bMCO-11 transfected V1 showed a long-lasting rise in bioluminescence signals. The time-lapse bioluminescence in VI upon white light visual stimulation in bMCO-11 expressing wild type mice is shown in Fig. 4E. Fig. 4F shows the kinetics of bioluminescence in V1 of bMCO-11 expressing wild type mice in response to repeated visual stimulation (indicated by the white bar, intensity near eye: 14 μW/mm^2^). In some instances, in bMCO-11 transfected mice, we observed visually evoked Ca^2+^-dependent bioluminescence signal propagation in the V1, which was detected by measuring the bioluminescence activity in different regions of the imaging window. The propagation and dynamics of the Ca^2+^-dependent bioluminescence activity in VI upon visual stimulation is summarized in Suppl. Fig. 4. Time-lapse bioluminescence mapping in V1 upon white light visual stimulation is shown in Suppl. Fig. 4A. Suppl. Fig. 4B shows the kinetics of the Ca^2+^-dependent bioluminescence signal in V1 in response to visual stimulation (intensity near eye: 14 μW/mm^2^).

**Fig. 4.**
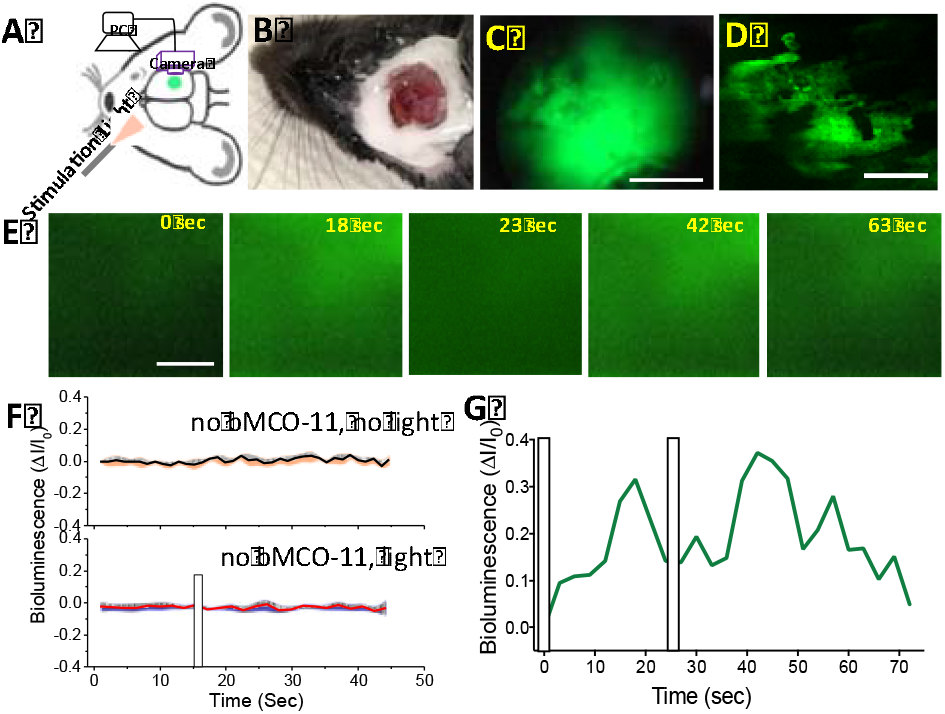
In-vivo monitoring of visually evoked bioluminescence activity in the visual cortex. (A) Schematic of the visual stimulation and bioluminescence imaging setup. (B) Optical window placed over visual cortex for bioluminescence imaging. (C) bMCO-11 expression monitored by epifluorescence imaging. Scale: 500 μm. (D) Zoomed confocal image of bMCO-11 expressing visual cortex. Scale bar: 50 μm. (E) Time-lapse bioluminescence images in V1 upon white light visual stimulation in the visual cortex of wild type mice transfected with bMCO-11. Scale: 200 μm. (F) Control study of Ca^2+^-dependent bioluminescence activity in wild type mice V1. Top: Bioluminescence of negative control (no bMCO-11 transfection and no visual stimulation light). Bottom: No change in Vl-bioiuminescence in negative control (no bMCO-11) in presence of visual stimulation (bar, intensity: 14 μW/mm¿). Average ± S.D. (G) Kinetics of Co^2+^-dependent bioìuminescence in VI in response to repeated visual stimulation (white bars, intensity: 14 μW/mm^2^)

### No change in cortical Ca^2+^-dependent bioluminescence activity in response to visual stimulation ofrdl mice

In order to evaluate the effect of retinal degeneration in correlation to the change in visually evoked activity of the visual cortex, time-lapse imaging of Ca^2+^-dependent bioluminescence signals in the primary visual cortex [VI] was carried out subsequent to visual stimulation. Since *rd1* mice are known to have fast and complete retinal degeneration, adult rdl and wild type mice with bMCO-11 transfected VI were used for comparative study. To confirm complete photoreceptor degeneration, SD-OCT imaging was performed. As shown in Fig. 5, in contrast to the wild type mouse retina (Fig. 5A), the *rd1* mouse retina [Fig. 5B] was found to lack the outer segment and photoreceptor layer. Kinetics of the Ca^2+^-dependent bioluminescence activity in VI neurons in response to visual stimulation (intensity: 20 μW/mm^2^) in *rd1* and wild type mice is shown in Fig. 5C. Upon visual stimulation, no significant rise in bioluminescence signal was observed in *rd1* mice. In contrast, wild type mice exhibited robust long-lasting Ca^2+^-dependent bioluminescence activity. Quantitative comparison of Ca^2+^-dependent bioluminescence activity in V1 with visual stimulation light ON and OFF for *rd1* and wild type mice are shown in Fig. 5D. As shown, a statistically significant difference in the change in Ca^2+^-dependent bioluminescence was observed between visual stimulation ON and OFF conditions for wild type mice.

**Fig. 5.**
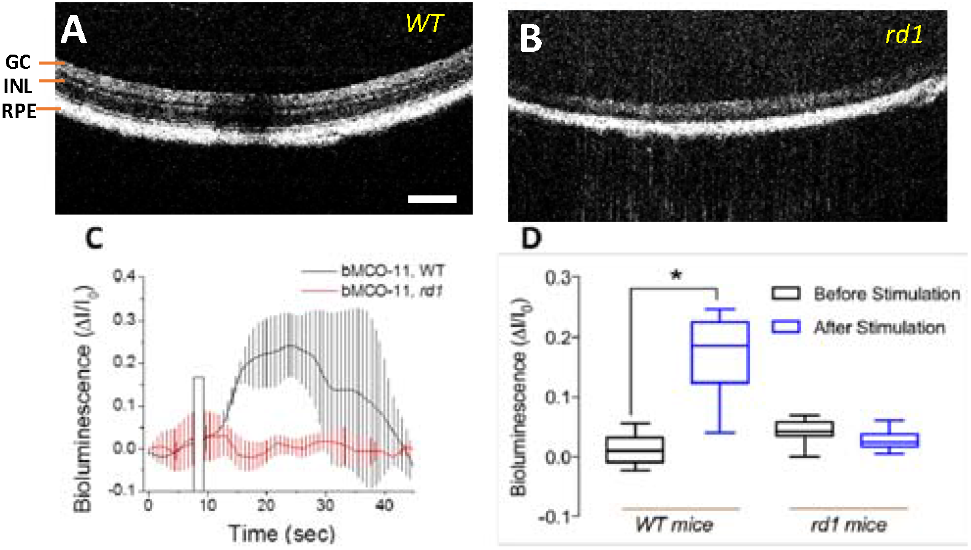
Quantitative comparison of visually evoked cortical activity monitored by bMCO-11 bioìuminescence between wild type and retinal degenerated mice. Representative SD-OCT image of wild type (A) and rd1 (B) mouse retinas. GC: Ganglion Cell layer; INL: Inner nuclear layer; RPE: Retinal pigment epithelium layer. Scale bar: 100 μm. (C) Kinetics of bioìuminescence cortical activity in VI in response to visual stimulation (intensity: 20 μW/mmĳ. The bar represents visual stimulation. (D) Quantitative comparison of bioluminescence cortical activity with visual stimulation light ON and OFF for rd1 and wild type mice.

## DISCUSSION

Current activity detection in diagnostic devices and sensory neural interfaces, including those in the visual cortex and retina, are either limited by spatial resolution [EEG, ERG etc] or invasiveness. Their use of penetrating electrodes can cause scar-tissue formation around the implanted area. Increasing the detection resolution requires higher electrode density and more current, thereby leading to heat production. Fluorescence-based neural activity monitoring, especially in visual systems, suffers from three major drawbacks: First, dependence on external excitation light prevents its use for study of neural activity in the retina, since the photoreceptors in the retina also get excited by the external fluorescence excitation light. Secondly, the use of light to excite fluorescence, especially for chronic stimulation, may substantially damage retinal and cortical neurons or any residual light-sensing function that might exist in the diseased or impaired retina. Additionally, using fluorescence excitation light and simultaneous optogenetic stimulation produces difficulties in decoupling optogenetic activation light from fluorescence excitation light. In order to achieve optical modulation of cellular activity and activity imaging, we have developed a chimera (bMCO-11) of ambient-light activatable opsin and Ca^2+^-sensitive bioluminescence protein. This allowed microscopic as well as mesoscopic bioluminescence imaging of cellular activation after localized single-cell as well as wide-area optical stimulation. The non-requirement of an external excitation light source for functional imaging (Ca^2+^-dependent bioluminescence) eliminates light-induced damage[26], thus allowing long-term activity assessment. Mesoscopic bioluminescence activity detection (in conjunction with stimulation) will provide a platform for screening the efficacy of therapeutic drugs in a high throughput manner.

Furthermore, the overlap between the activation spectrum of the opsin-domain [MC0-11] and the emitted light spectrum from the GeNL domain in bMCO-11 chimera leads to a reduced requirement for external optogenetic stimulation light. Compared with other strategies, an optogenetic method is cell-specific and provides higher resolution. The optogenetic stimulation method has been used for modulation of neural activity for therapeutic usage[27] including vision restoration through stimulation of the retina[28-31]. Similar to patterned electrical stimulation of the visual cortex (in the case of optic nerve damage or enucleation), optogenetics also provides the possibility for high-resolution stimulation. Our optogenetic studies in retinal degenerated mouse models suggest that a photoreceptor degenerated retina can be made highly photosensitive, enabling vision restoration at low light levels. Though the primary visual cortex is known to maintain its retinotopy in subjects with retinal degeneration despite prolonged visual loss, detailed knowledge of how optogenetic sensitization of higher order neurons manifests in the restoration of visual cortical activity is currently lacking. Thus, there is a need for mapping changes in the visual cortical activity as progression of retinal degeneration and subsequent vision restoration by optogenetic sensitization of the retina occurs. Using bMCO-11, we characterized visually evoked activity in retinal degenerated as well as wild type mice. While no change in Ca^2+^-dependent bioluminescence signal was observed in *rd1* mice, robust increase in bioluminescence was noted in the primary visual cortex of wild type mice upon visual stimulation. Using bandwidth engineering, we have synthesized bioluminescent MC0 [bMCO-11] with an overlapping activation spectrum that led to perpetuation of the bioluminescence response. Similarly, an inhibitory opsin [32] with an overlapping activation spectrum can be integrated with the bioluminescence [GeNL] domain. The passive modulation of cellular activity will lead to modulation of neurological functions. For example, already-activated, hyper-sensitized pain neurons [27] expressing bMCO will generate bioluminescence light, which will up-regulate (in case of excitatory opsins as used in the present study) or down regulate the activity of the targeted cells expressing bMCO containing inhibitory opsins leading to pain modulation.

The use of completely separated activation spectra for the opsin and the emitted bioluminescence in the neuronal activity sensor would allow bidirectional optogenetic control of cellular activity triggered by light activation of the opsin (write) and monitoring of signal propagation in a neural network (read) visualized by bioluminescence imaging. This will lead to the development of a modular and scalable interface system with the capability to serve a multiplicity of applications to modulate and monitor large-scale activity in the nervous system. Integration of a voltage-sensitive bioluminescent sensor with the multicharacteristic opsin will enable us to modulate and image neural activity in the retina and visual cortex with high temporal and spatial resolution, providing details about disease progression and restoration. The MCO component can be activated using GeNL bioluminescence and vice versa. As the bioluminescence signal is weak, the cyclic activation of the MCO moiety will serve to amplify the signal and therby enable long neural activity imaging. By combining the complementary advantages of both chemical and optical methods, the same genetic construct can be used to control neuronal activity via both chemical and optical stimuli. These features make bMCO-11 useful for the comprehensive interrogation of neuronal circuits. In the long term, we aim to demonstrate a clinically viable optical stimulation device that can reliably provide stimulation input to the visual cortex and simultaneous recording of cortical activity.

## CONCLUSIONS

We successfully demonstrated the development and use of bioluminescent multi-characteristic opsin (bMCO-11) for efficient activation and excitation-free recording of cellular activity on a microscopic to mesoscopic scale. Cyclic activation of the opsin-domain in the bMCO-11 expressing cortical neurons by visually evoked bioluminescence led to persistent Ca^2+^ influx. This allowed us to continuously monitor visual cortical activity in wild type and retinal degenerated mice without requiring an additional external excitation source. Simultaneous bioluminescence measurements of cortical activity in response to visual stimulation presents a new modality to efficiently monitor the visual function longitudinally during disease progression and therapeutic treatments. Advancements in temporal sensitivity of the bioluminescent-sensors and spectrally separated activation of narrow band actuators will lead to development of a modular and scalable interface system with the capability to serve a multiplicity of applications to monitor and modulate large-scale activity in the nervous system.

## MATERIAL AND METHODS

### Construction and characterization of Bioluminescent Multi-Characteristic Opsin

We synthesized a chimeric bioluminescent multi-characteristic opsin (bMCO-11) by the fusion of light-activatable domain (MCO-11) and Ca^2+^ sensing bioluminescent domain (GeNL) using DNA synthesizer. CAG promoter was used upstream of light-activatable domain to express in HEK cells as well as cortical neurons. CAG is known to preferentially transfect excitatory neurons in all layers of the visual cortex. The fusion plasmid sequence was verified and cloned into pET-vector using the restriction enzymes XhoI and BamHI. Fig. 1A shows the Domain architecture of bMCO-11. The plasmid containing bMCO-11 was lyophilized and dissolved in sterile TE buffer before transfection. DNA gel electrophoresis was carried out to verify the size and purity of the bMCO-11 gene (digested by restriction enzymes BamHI and Sail with restriction fragments).

### Theoretical modeling of Bioluminescent MultiCharacteristic Opsin

The theoretical modeling of bMCO-11 was carried out using web-based protocol, “RaptorX”. Further, the secondary structure was predicted by OCTOPUS, which uses a combination of hidden Markov models and artificial neural networks. First, homology search using BLAST is performed to create a sequence profile, which is used as the input to a set of neural networks that predict both the preference for each residue to be located in a transmembrane (TM), interface (I), close loop (L) or globular loop (G) environment. The preference for each residue to be inside (i) or outside (o) of the membrane is also predicted. In the final step, these predictions are used as input to a two-track hidden Markov model to calculate the most likely topology.

### Cell culture and bMCO-11 plasmid transfection

HEK293 cells were cultured in Petri dishes and maintained in DMEM/F-12 with 10% fetal bovine serum, and 0.05mg/mL gentamycin. The cultures were maintained at 37 °C in a 5% CO_2_ humidified atmosphere. The cells were transfected with bMCO-11 plasmids using Lipofectamine 3000 [Invitrogen]. 2 days after transfection, visualization of the fluorescence was carried out under an epifluorescence microscope.

### Fiber-optic optogenetic stimulation

A single mode optical fiber coupled to a supercontinuum laser source (NKT Photonics) delivered the light to the cells (on a petridish) for optogenetic stimulation. The fiber was mounted, and μ the petri dish for wide area stimulation. The divergent light from the fiber could cover ~20 mm diameter at the sample plane so that the patched cells could be homogenously stimulated in the dish. A power meter (Newport 1815-C) was used to quantify the light intensity at the sample plane. The pulse width was controlled by an electro-mechanical shutter, synchronized with the electrophysiology recording system [Molecular Devices].

### Patch-clamp recording setup

Cells, transfected with bMCO-11, were incubated with all-trans retinal (ATR, 1 μM) for 6 hours before conducting the patch-clamp experiments. Just before patch clamp measurements, coelenterazine-h (10 ¼M) was added. Inward photocurrents rents in bMCO-11 transfected cells were recorded using the patch clamp. The patch-clamp recording setup consists of an inverted Nikon fluorescence microscope platform using an amplifier system (Axon MultiClamp 700A, Molecular Devices). Visualization of the bMCO-11 fluorescence in transfected cells was carried out under epifluorescence and was dark adapted thereafter for >5 min. Micropipettes were pulled using a two-stage pipette puller to attain resistance of 3 to 5 MΩ when filled with intra-cellular solution. The micropipette was filled with a solution containing (in mM) 130 K-Gluoconate, 7 KC1, 2 NaCl, 1 MgCl2, 0.4 EGTA, 10 HEPES, 2 ATP-Mg, 0.3 GTP-Tris and 20 sucrose. The micropipette-electrode was mounted on a micromanipulator. The extracellular solution contained (in mM): 150 NaCl, 10 Glucose, 5 KC1, 2 CaCl2, 1 MgCl2 was buffered with 10 mM HEPES (pH 7.3). Series resistance was adjusted in the amplifier using the inbuilt Rs compensation control provided in the V-Clamp pane of MultiClamp 700A. Photocurrents were measured while holding cells in voltage clamp at −70 mV. The electrophysiological signals from the amplifier were digitized using Digidata 1440 (Molecular devices), interfaced with patch-clamp software (pClamp, Molecular Devices). pClamp 10 software was used for data analysis.

### Single-cell targeted microscopic optical stimulation and bioìuminescence activity detection

Cells, transfected with bMCO-11, were incubated with all-trans retinal (ATR, 1 μM) for 6 hours before conducting the single-cell targeted optical stimulation and bioìuminescence activity detection using upright confocal laser microscopy (Olympus Fluoview 1000). Coelenterazine-h (10 μM) was added to the cells before activity measurements. Visualization of the bMCO-11 fluorescence in transfected cells was carried out under confocal fluorescence (excitation: 488 nm, emission: 510-550 nm) and was dark adapted thereafter for >5 min before optical stimulation and bioluminescence activity measurements. The petridish and objective was enclosed in a light-tight box and room light was turned off. After identification of the transfected cell(s), the cell (region) of interest was selected and scanned using focused 488 nm activation laser microbeam. Immediately after switching off the laser activation, bioluminescence image was acquired using confocal PMT-detection by scanning wider region of interest (including the stimulated cell as well as the non-targeted cell).

### Wide-area optogenetic stimulation and bioluminescence detection of cellular activity

For conducting the wide-area optical stimulation and bioluminescence activity detection of cellular activity, cells transfected with bMCO-11 were incubated with all-trans retinal (ATR, 1 μM) for 6 hours before measurements. Cells in petridish was placed in an inverted automated epifluorescence microscope (Discovery, Molecular devices) and covered with a dark enclosure and room light was turned off. Visualization of the bMCO-11 fluorescence in transfected cells was carried out under epifluorescence (excitation: 480-500 nm, emission: 520-550 nm) and was dark adapted thereafter for >5 min before optical stimulation and bioluminescence activity measurements. The petridish and objective was enclosed the cells before activity measurements. For wide-area optical activation, cells were irradiated with a narrow band light (480-500 nm) from the Xenon lamp delivered via a dichroic mirror and microscope objective (20X]. The pulse width of the optical activation was controlled by an electro-mechanical shutter (Molecular Devices]. Time-lapse bioluminescence images were acquired via band pass filter (510-550 nm) using cooled Photometrics camera.

### Mouse preparation

Retinal degenerated mice (B6.CXB1-Pde6b^rd1^/J) and wild type (C57BL/6J) were obtained from Jackson laboratory and bred in the animal facilities of the Nanoscope Technologies. Mice were maintained on a 12:12 light cycle. The mice were treated humanely in strict compliance with IACUC on the use of animals in research.

### Spectral Domain Optical Coherence Tomographic (SDOCT) imaging of retina

To confirm complete photoreceptor degeneration in the rd1 mice, the animals were anesthetized with a mixture of ketamine (65 mg/kg], xylazine (7.5 mg/kg], and acepromazine (0.5 mg/kg) and mounted on a maneuverable imaging platform. 1~2 drops of 1% tropicamide was topically applied to eyes for pupil dilation. The cornea was kept moist with a balanced salt solution during the measurement period. Spectral Domain-OCT (SD-OCT) (33-35) imaging of retina was performed in *rd1* as well as wild type mice.

### Cortical transfection of bMCO-11 plasmids, optical window placement for bioluminescence imaging

Craniotomy was carried out on >12 weeks old anesthetized rd1 and wild type mice, over the primary visual cortex. After removal of the dura mater, solution of bMCO-11 plasmids in Jetprime transfection reagent (Polyplus) was injected at multiple locations of the visual cortex using a microneedle-syringe. The optical window (5 mm glass coverslip) was placed over the visual cortex for bioluminescence monitoring and glued to the skull using dental cement.

### Evaluation of bMCO-11 expression in the primary visual cortex using confocal microscopy

For evaluation of bMCO-11 expression in the visual cortex, after anesthetization of the control (non-injected) and bMCO-11 injected mice, the rd1 or wild type mice were placed under upright confocal laser microscopy (Olympus Fluoview 1000]. 4X microscope objective and camera was used for in-vivo wide area epifluorescence imaging excitation: 488 nm, emission: 510-550 nm]. 10X objective was used to the visualize expression in cortical neurons via three-dimensional confocal fluorescence scanning through the optical window placed over the visual cortex.

### Measurement of Bioluminescent visual cortical activity upon visual stimulation

1 week after cortical transfection, the animals were anesthetized with a mixture of ketamine (65 mg/kg], xylazine (7.5 mg/kg], and acepromazine (0.5 mg/kg) and mounted on a maneuverable imaging platform. 1~2 drops of 1% tropicamide was topically applied to eyes for pupil dilation. The cornea was kept moist with a balanced salt solution during the measurement period. For visual stimulation white LED light was placed in front of the eye. The intensity near the eye was measured using a power meter (Newport]. A CMOS sensor (OnSemi MT9P031) and imaging lens was mounted over the optical window to monitor bioluminescence activity of visual cortex with and without visual stimulation. The imaging lens was adjusted to focus the top layer of visual cortex on to the camera. Coelenterazine-h (200 mg/kg body-weight) or Furimazine (200 mg/kg bodyweight) was delivered via intraperitoneal injection (final volume: 500 μl) 10 minute before bioluminescence imaging. The gain and exposure time of the camera was kept fixed and time-lapse (~ 2sec) images were acquired before and after visual stimulation. The room light was turned off and monitor was kept at lowest brightness for minimizing background noise during bioluminescence measurements.

## AUTHOR INFORMATION

### Author Contributions

The manuscript was written through contributions of all authors. / All authors have given approval to the final version of the manuscript.

## ACKNOWLEDGMENT

The authors would like to thank the National Eye Institute grants, 1R01EY02821601A1, 2R44EY02590502A1, 1R43EY025905-01, 1R43EY026483-01, 3R43EY025905-01S1,1R01EY025717-01A1 and JST SENTAN.

## Supplementary Information

**Suppl. Fig. 1.**
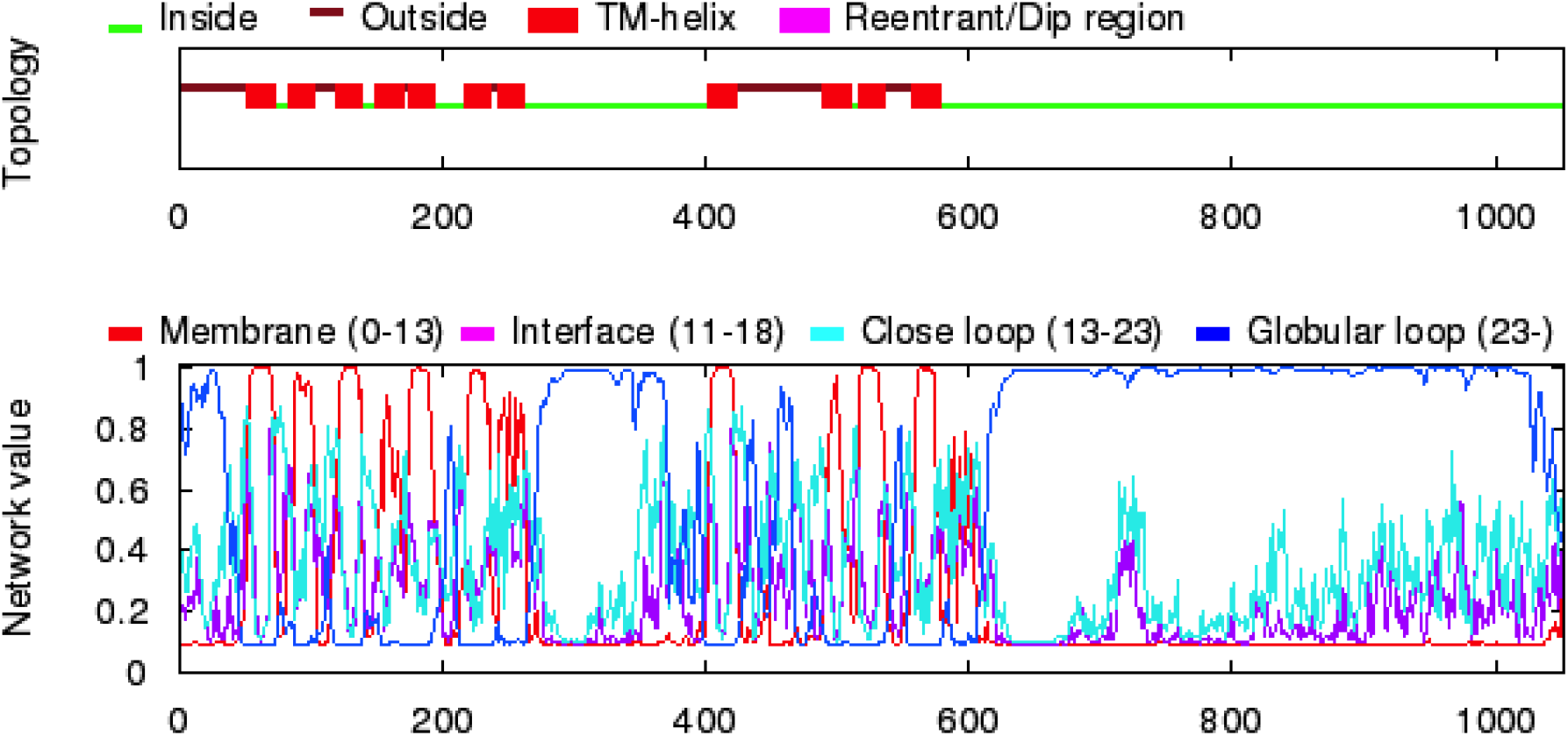
Theoretical modeling of the three-dimensional arrangement of amino acid chains of bMCO-11. The topology of trans membrane helices in bMCO-11 is predicted using the software “OCTOPUS” which uses improved topology prediction by two-track ANN-based preference scores. X-axis represents the position of amino acids and Y-axis represents the theoretically calculated network value.

**Suppl. Fig. 2.**
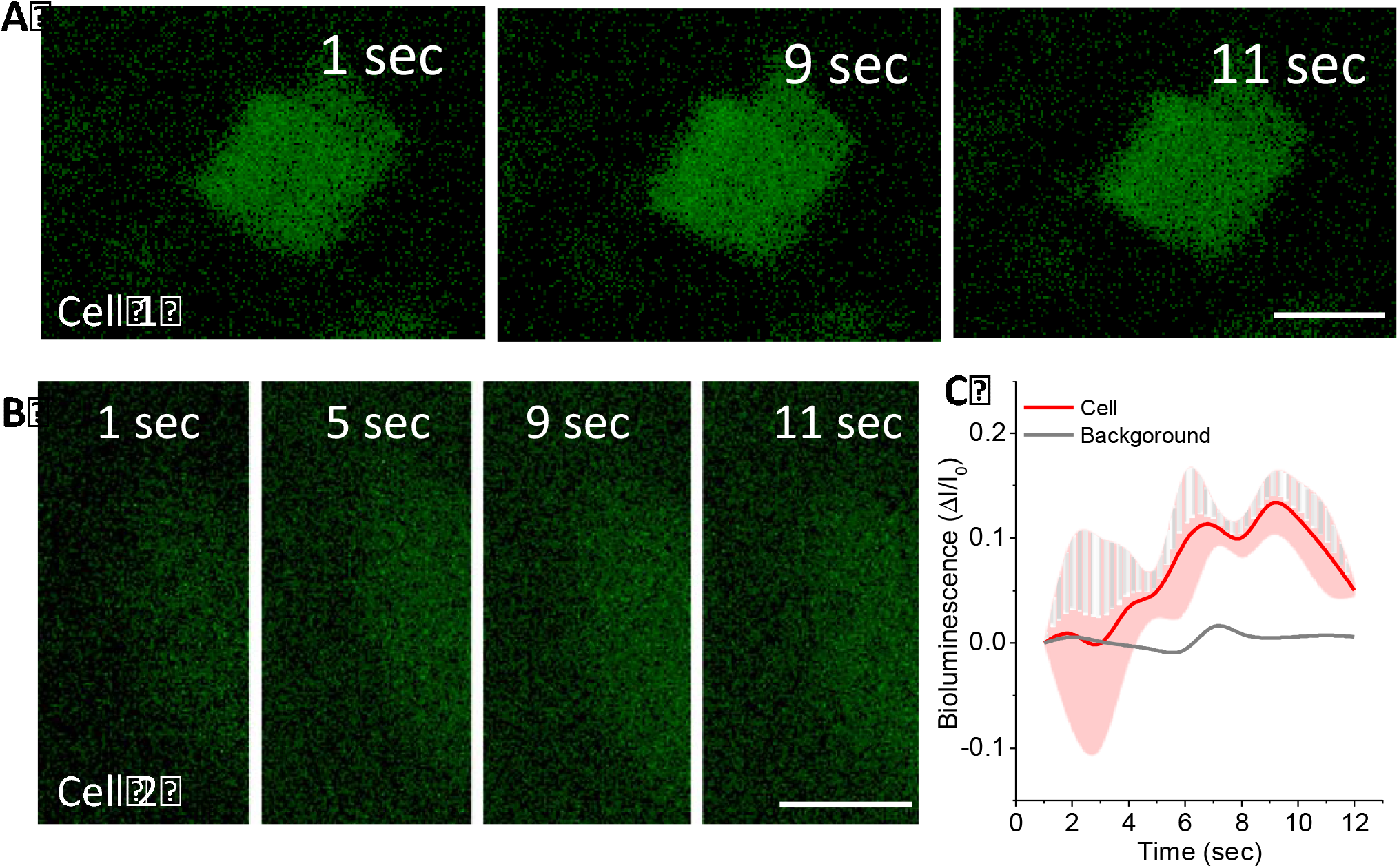
Monitoring single cell Ca^2+^-dependent bioluminescence after localized optical stimulation. (A, B) Time-lapse bioluminescence images of HEK cells upon localized stimulation. Scale: 10 μm. (C) Kinetics of Ca^2+^-dependent bioluminescence changes subsequent to localized optical stimulation of single cell.

**Suppl. Fig. 3.**
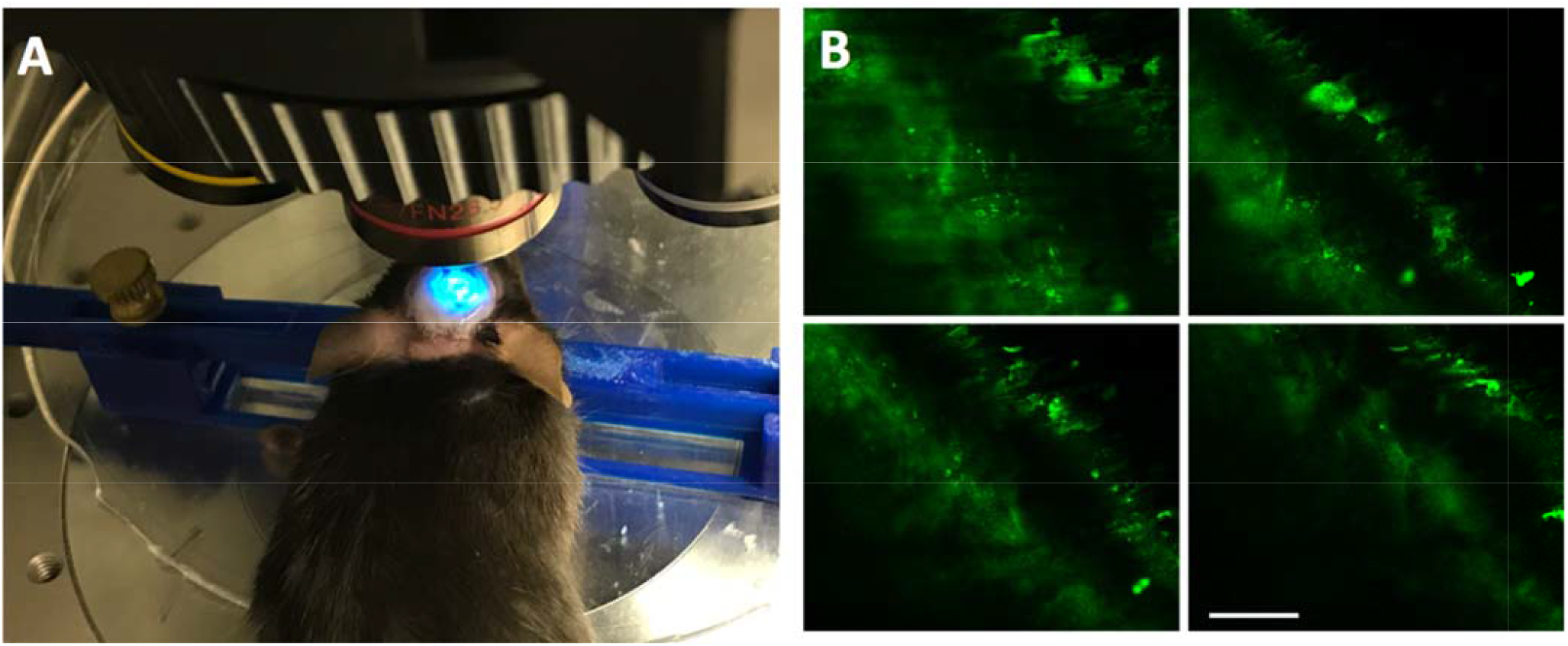
Monitoring bMCO-11 expression in visual cortex using confocal microscopy. (A) Confocal imaging platform. (B) Serial-sectioned confocal image showing bMCO-11 expression in visual cortex. Scale: 200 μm.

**Suppl. Fig. 4.**
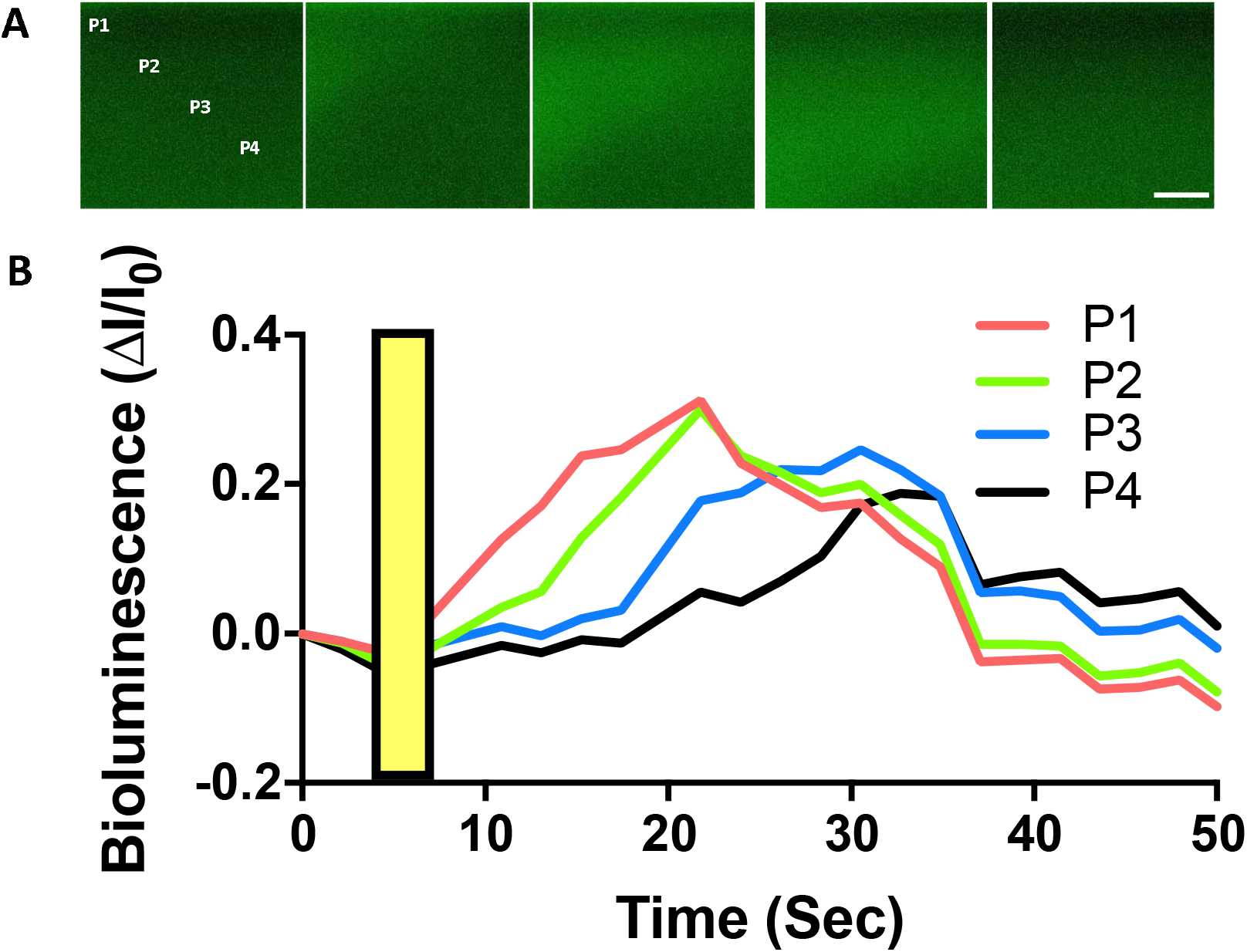
Dynamics of visually evoked Ca^2+^-dependent bioluminescence activity. (A)Time-lapse bioluminescence in V1 upon white light visual stimulation. Scale: 200 μm. (B) Kinetics of bioluminescence in V1 in response to visual stimulation (orange bars, intensity: 14 μW/mm^2^). The bioluminescence intensity was monitored at different locations (P1, P2, P3, P4) over time. The temporal evolution of bioluminescence signal varies at different locations due to the propagation of the bioluminescence signal associated with cortical activity.

